# Low noise HRTFs and delay line corrections are detrimental to the prediction of ITD discrimination thresholds from environmental statistics

**DOI:** 10.1101/2022.09.09.507313

**Authors:** Maria Juliana Gutiérrez Camperos, Thaís Caroline Gonçalves, Bóris Marin, Rodrigo Pavão

## Abstract

Interaural Time Difference (ITD) is the main cue for azimuthal auditory perception in humans. ITDs at each frequency contribute differently to azimuth discrimination, which can be quantified by their azimuthal Fisher Information. Consistently, human ITD discrimination thresholds are predicted by the azimuthal information. However, this prediction is poor for frequencies below 500 Hz. Such poor prediction could be ascribed to the strategy of quantifying azimuthal information using HRTFs obtained in unnaturalistic anechoic chambers or by using a direct method which does not incorporate the delay lines proposed by the Jeffress-Colburn model. In the present study, we obtained ITD discrimination thresholds from extensive sampling across frequency and ITD, and applied multiple strategies for quantifying azimuthal information. These strategies employed HRTFs obtained in realistic and anechoic chambers, with and without considering delay lines. We found that ITD discriminability thresholds across the complete range of frequencies are better predicted by azimuthal information conveyed by ITD cues when (1) we use naturalistic high-noise HRTFs, and (2) ITD delay compensation is not applied. Our results support that auditory perception is shaped by natural environments, which include high reverberation in low frequencies. Moreover, we also suggest that delay lines are not a crucial feature for determining ITD discrimination thresholds in the human auditory system.

**GRAPHICAL ABSTRACT:** 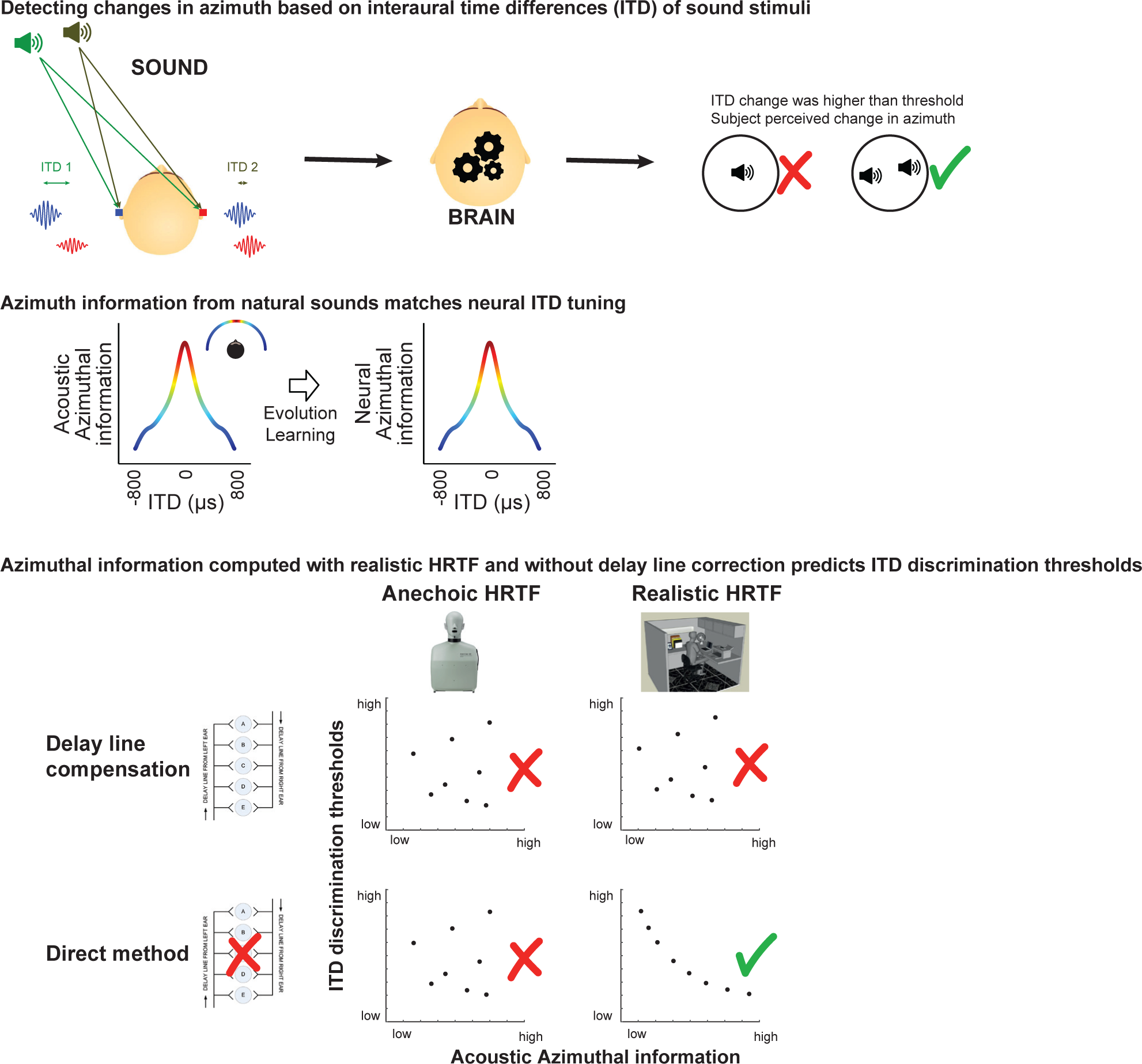

## INTRODUCTION

Humans localize low-frequency sounds along the horizontal plane based mainly on the difference between the time of arrival of the acoustic signal at the left and right ear, the so called ITD (Interaural Time Difference) (Strutt, 1907; Middlebrooks and Green 1991). Sound emanating from a given azimuth reaches the ears and is decomposed across frequency bands by the cochleae. The spatial percept is computed from the ITD cues by auditory circuits in the brainstem (Schnupp, Nelken, and King 2011).

ITD signals for a given auditory stimulus can be estimated from head-related transfer functions (HRTF) and frequency filters of a cochlear model (**Fig. 1A**). The frequency components within the critical band of the cochlear filter interfere and generate the instantaneous ITD time series, which fluctuates around a given mean (Pavão et al. 2020). **Figure 1B** shows the mean and standard deviation of ITD as a function of azimuth, computed across the low-frequency range.

**Figure 1:**
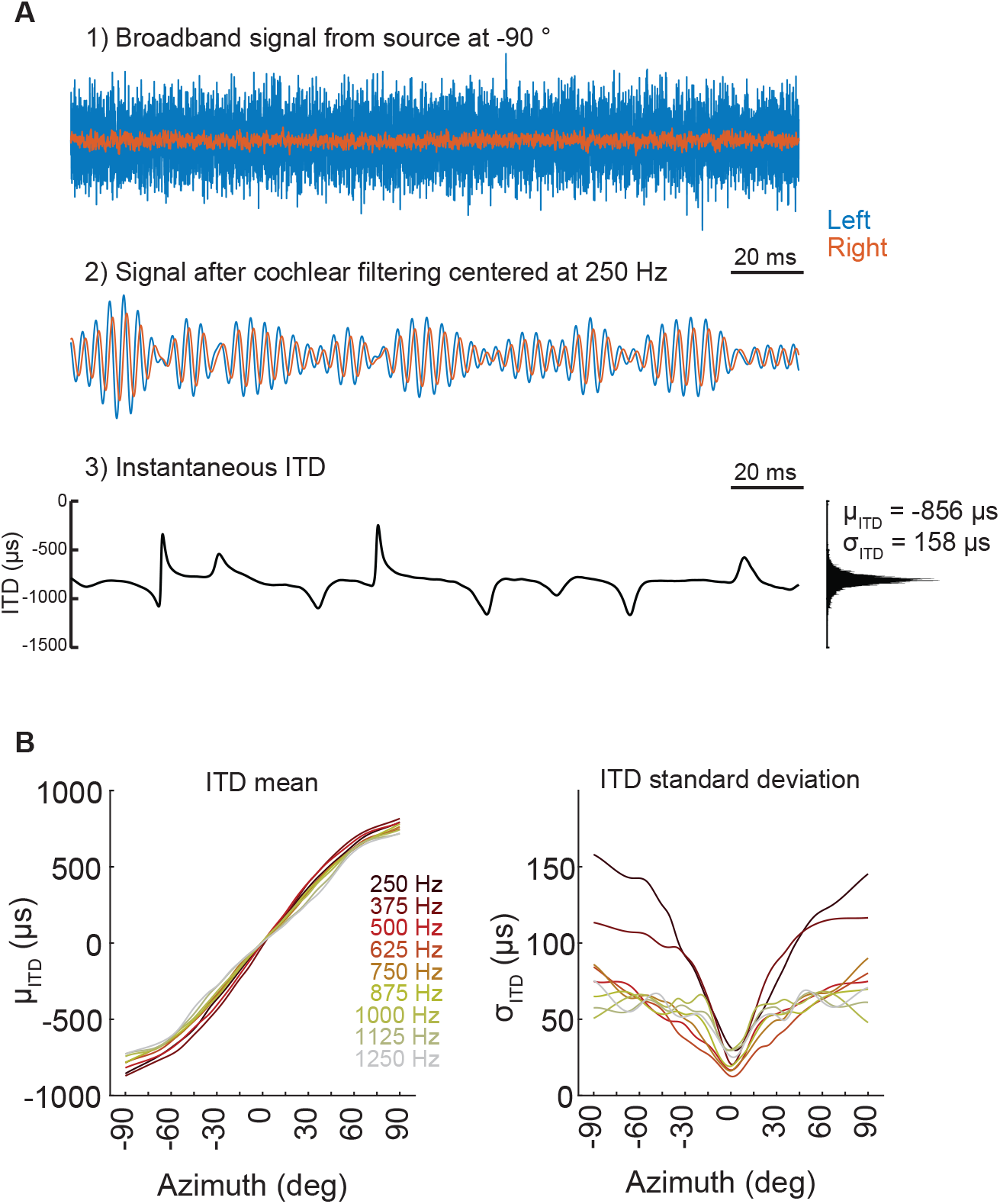
Computing ITD mean and standard deviation across azimuth and frequency. (**A**) 1) Example of white noise broadband signal convolved with human head-related impulse responses (HRIRs; speaker at −90° azimuth). Blue and red curves depict the resulting signal for left and right ears. 2) Signal obtained after band-pass filtering by cochlear model (centered at frequency of 250 Hz). 3) Left: the black curve depicts instantaneous interaural time difference (ITD; computed from instantaneous phase difference between left and right channels). Right: histogram of the distribution of instantaneous ITD, for the −90° azimuth at 250 Hz, with the corresponding ITD mean (μ_ITD_) and standard deviation (σ_ITD_). (**B**) Cubic spline interpolation of μ_ITD_ vs azimuth (left) and σ_ITD_ vs azimuth (right) for each frequency, computed using the CIPIC HRTF database (Algazi et al. 2001).

The mean instantaneous ITD (μ_ITD_) and its standard deviation (σ_ITD_) across azimuth influences the amount of azimuthal information conveyed by ITD cues, which can be estimated using Fisher information. Higher Fisher information (FI_ITD_) in **Figure 2** indicates smaller decoding errors when estimating azimuth from ITD cues: this corresponds to azimuth ranges for which the μ_ITD_ curve (**Fig. 1B**, left) is steeper and σ_ITD_ is lower (**Fig. 1B**, right). In (Pavão et al. 2020), FI_ITD_ was succesfully employed to predict not only neural activity in rodent brainstem (Pecka et al. 2008; McAlpine, Jiang, and Palmer 2001) but also ITD discrimination thresholds obtained from psychophysics (Pavão et al. 2020; Mills 1958).

**Figure 2:**
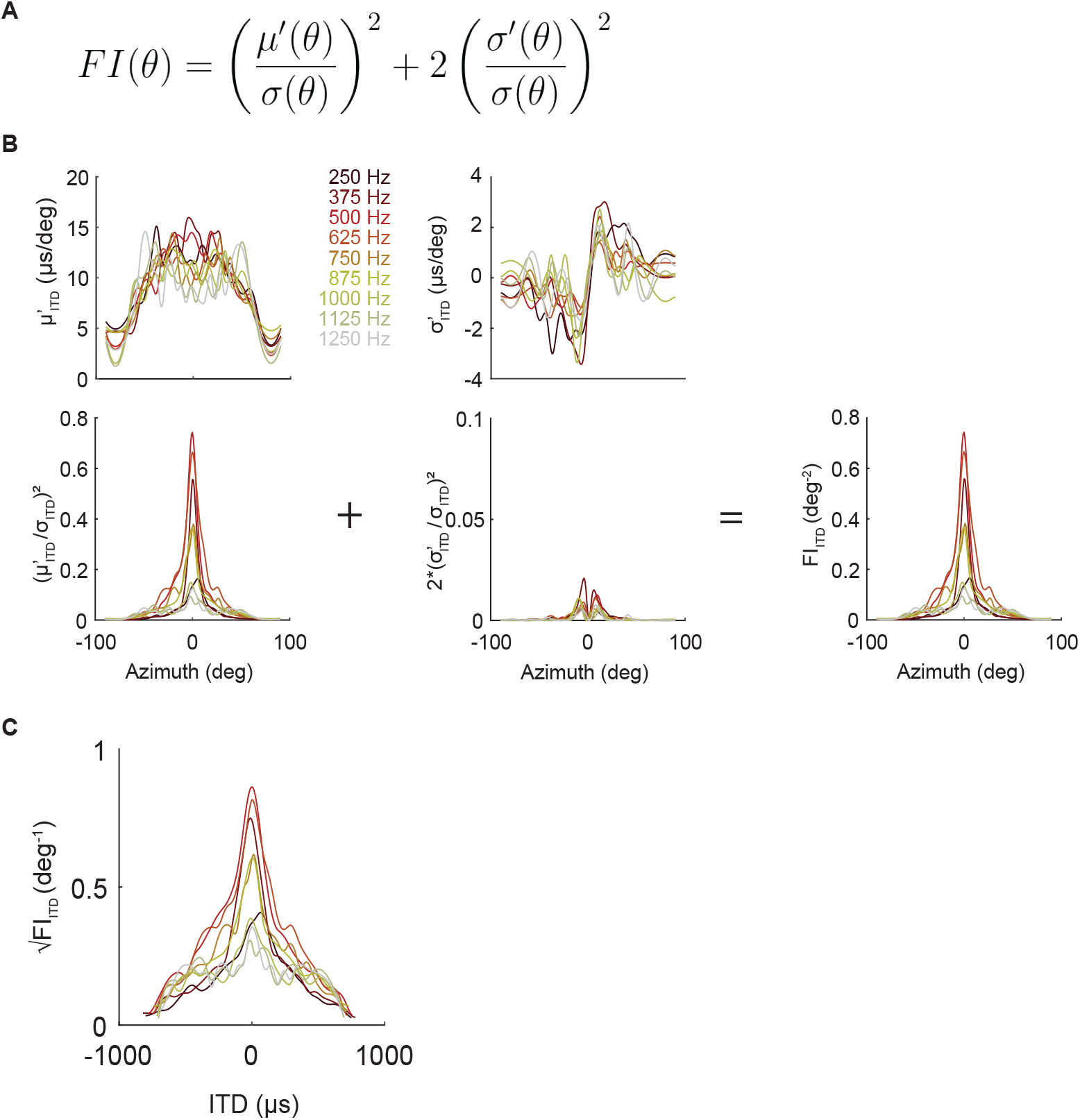
Computing ITD Fisher information (FI_ITD_). (**A**) Assuming that the conditional distributions of instantaneous ITD given azimuth *p(ITD*|*θ)* are Gaussian, the Fisher information (FI) is computed from the derivative of their mean μ’(θ), the derivative of the standard deviation σ’(θ) and the standard deviation itself σ(θ). When applied to the context of sound localization, FI_ITD_ reflects the amount of azimuthal information carried by environmental ITD cues. (**B**) Top row: μ’_ITD_(θ) (derivative of mean ITD as a function of azimuth; left) and σ’_ITD_(θ) (derivative of ITD standard deviation as a function of azimuth; right). Bottom row: the sum of the first and second terms (left and center plots) of equation in A results in FI_ITD_ across azimuth (right). Notice that FI_ITD_ (bottom right) is dominated by the first term of the equation (bottom left). (**C**) Square root of FI_ITD_ across ITD. Discrimination thresholds are shown to be proportional to √FI_ITD_. Azimuths have been replaced with the corresponding μ_ITD_ (Fig. 1B, left).

ITD discrimination thresholds change across frequency and azimuth (Mills 1958; Yost 1974; Brughera, Dunai, and Hartmann 2013): highest ITD discriminability is found near 750 Hz close to the midline. In addition, (Pavão et al. 2020) showed that √FI_ITD_ predicts ITD discriminability from 500 to 1250 Hz, but not for 250 Hz. Our premise is that estimating √FI_ITD_ through methods which mimic auditory mechanisms and reflect properties of natural environments could improve prediction of ITD discriminability. The direct method for estimating √FI_ITD_ described in (Pavão et al. 2020) was based on a single HRTF database for which, √FI_ITD_ curves are asymmetric, and ITDs were computed from the acoustic signals independently of their amplitude and delay compensation. These are potential caveats given that human head and ears are symmetrical, there is a sound pressure threshold for exciting the basilar membrane, and whether delay lines exist in humans or not is still under debate. In the present study, we investigate different methods for estimating FI_ITD_ that deal with these potential issues. We found that the direct method applied to the CIPIC HRTF database (which has higher σ_ITD_ for low frequencies) led to the highest accuracy in predicting ITD discriminability thresholds.

## RESULTS

Our main results consist in (a) contrasting different methods for estimating √FI_ITD_ (check Materials and Methods section for details) (**Figure 3**), (b) estimating ITD discriminability (dITD) thresholds in human subjects (**Figure 4**) and (c) predicting dITD thresholds using different √FI_ITD_ estimates (**Figure 5**).

**Figure 3:**
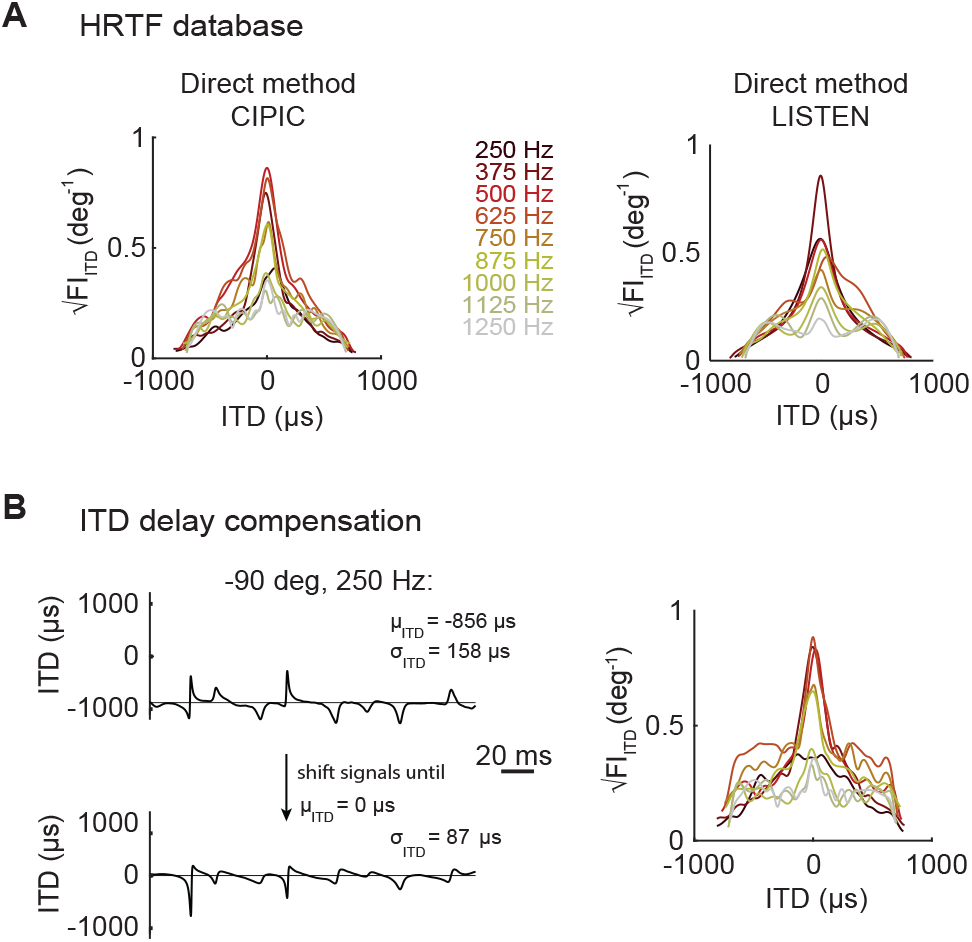
Methods for computing ITD Fisher Information and resulting√FI_ITD_. √FI_ITD_. Each method contemplates different strategies of processing ITD (left column) and the obtained √FI_ITD_ across frequency and azimuth (replaced with the corresponding μ_ITD_) (right column). (**A**) Direct method: we considered variations in the HRIR by taking the CIPIC (left) and the LISTEN HRTF databases (right). (**B**) Compensating ITD delay by the estimated μ_ITD_, which synchronizes low amplitude components of the left and right signals, potentially reducing σ_ITD_. This panel depicts results for the CIPIC HRTF database.

**Figure 4:**
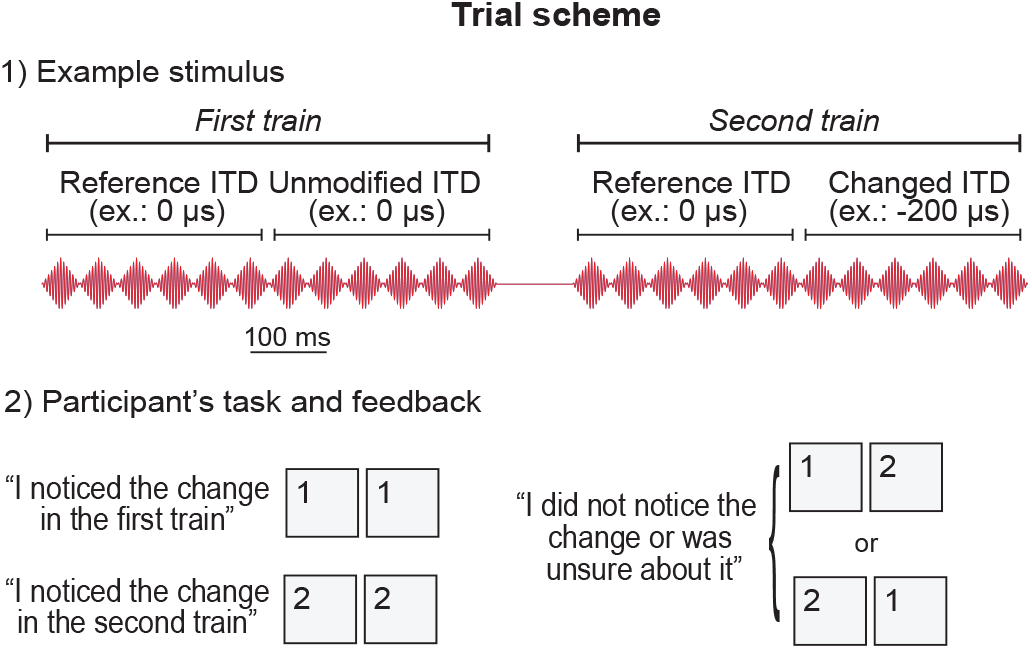
Experimental design for ITD discrimination task. Participants listened to a pair of trains of beeps and responded in which one they perceived a change in ITD, by pressing buttons “1” or “2”.

**Figure 5:**
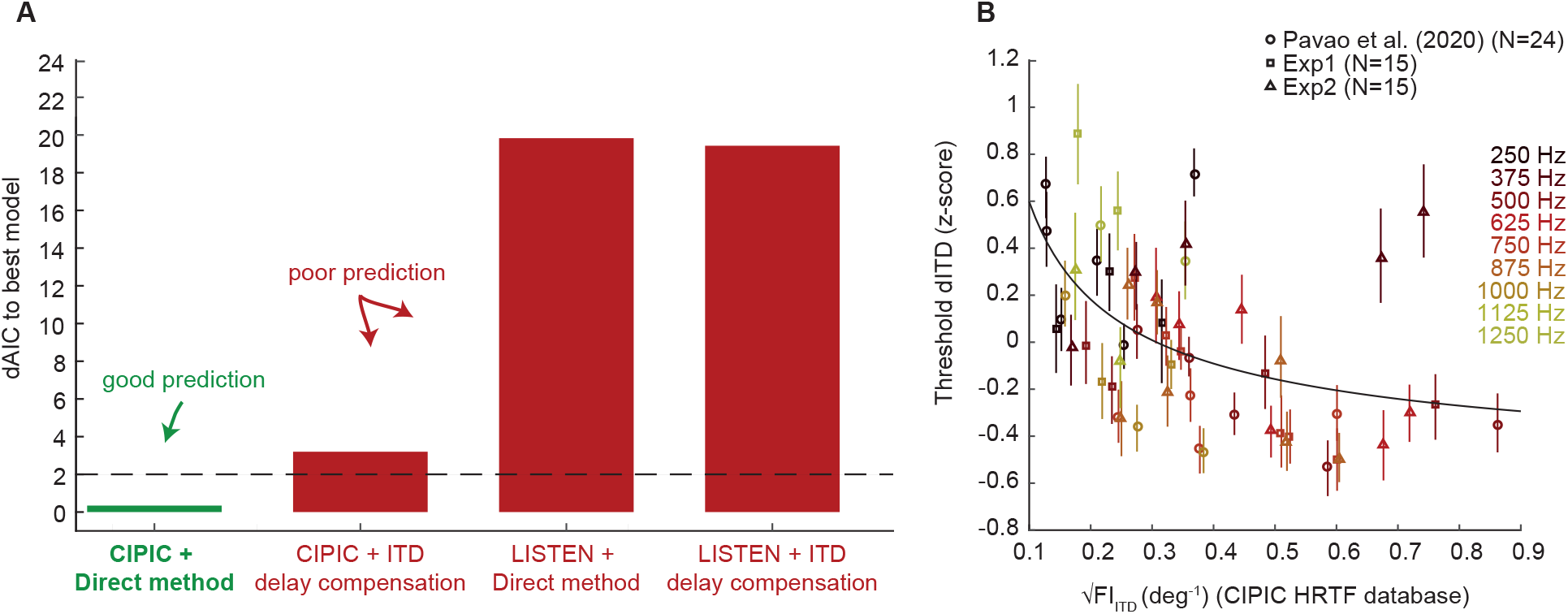
Prediction of dITD thresholds by √FI_ITD_. (**A**) Models with dAIC values lower than 2 are considered the best. The green and red bars show dAIC obtained from CIPIC and LISTEN HRTF databases using the direct method and delay compensation. The direct method applied to the CIPIC database leads to the best prediction of dITD thresholds. (**B**) Joint data from the experiments shown in Figure 4 supplement as a function of √FI_ITD_ estimates using the direct method based on CIPIC.

### Contrasting different methods for estimating √FI_ITD_

All different methods led to similar √FI_ITD_ distributions, with high information for ITDs corresponding to frontal azimuth across frequency (**Figure 3** and **Figure 3 supplement**). The most pronounced difference arose from methods that considered different HRTF databases, for frequencies below 500 Hz: √FI_ITD_ obtained using the LISTEN database were systematically higher than those calculated from the CIPIC database (**Figure 3A**). This could indicate that the anechoic chamber used to construct LISTEN is more effective in reducing low-frequency reverberations.

Another significant difference emerged for methods that considered ITD delay compensation (**Figure 3B**, right): √FI_ITD_ estimations in the periphery were higher than those obtained by the direct method (without compensation, **Figure 3A**, top right).

### Human dITD thresholds

In order to estimate dITD thresholds, we applied a psychophysics protocol in which the subjects had to detect ITD changes in the first or second train of beeps presented in the trial (**Figure 4**). ITD changes ranged from 1 to 200 μs from the reference. For each condition (combination of frequency and reference ITD), the estimated dITD threshold was the ITD value which maximally separated the subject’s correct and incorrect/unsure responses. This is essentially the same protocol as the one described in (Pavão et al. 2020), but applied to additional subjects (Experiments 1 and 2). **Figure 4 supplement** shows dITD thresholds averaged across subjects for each condition from the three datasets/experiments, which differ by the set of conditions presented to the independent pool of subjects. There is no clear correlation between dITD threshold and reference ITD or frequency – the only cues present in the stimuli. These considerations motivated investigating predictors based on natural statistics.

**Figure 3 supplement:**
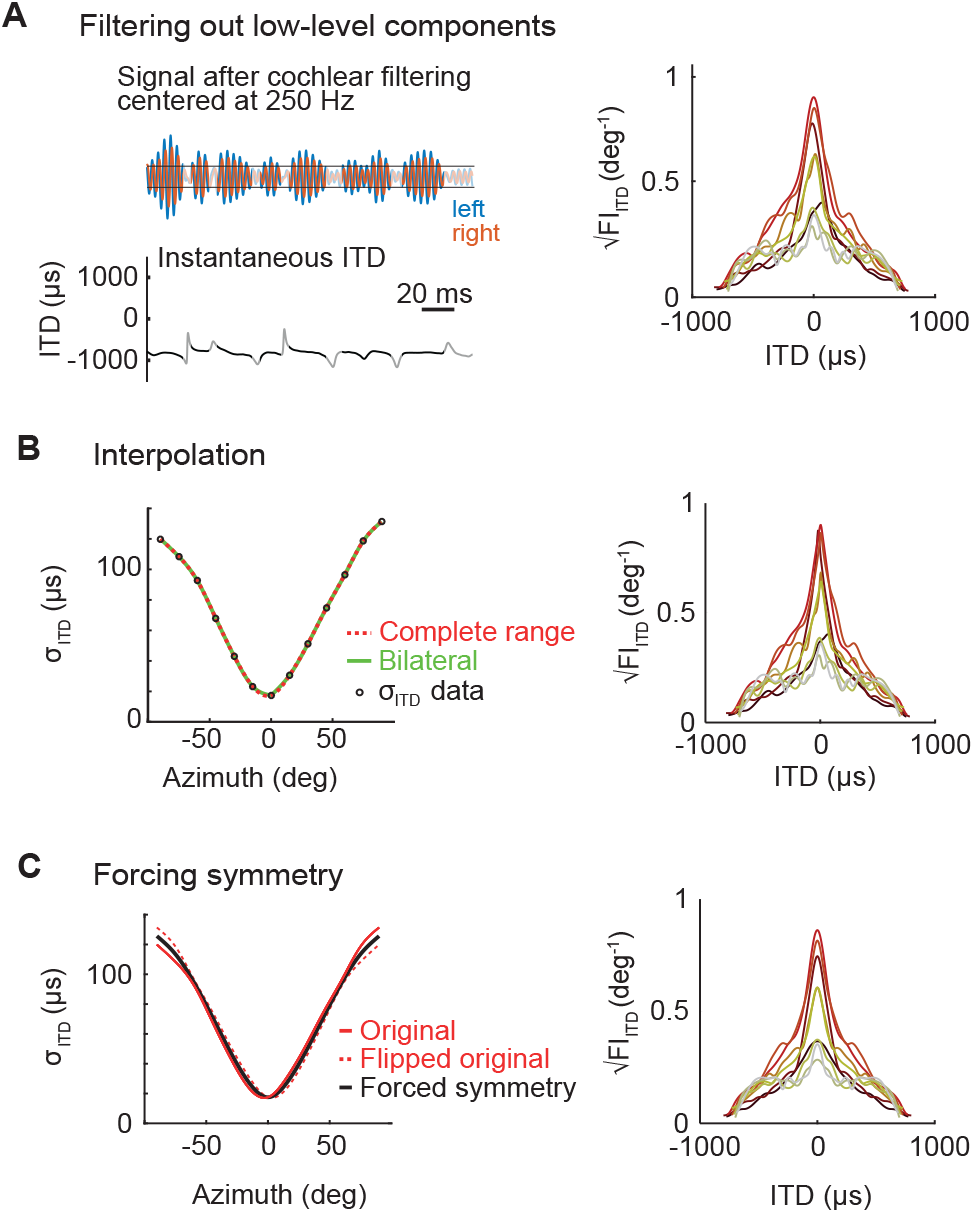
Additional methods for computing ITD Fisher Information and resulting √FI_ITD_. (**A**) Filtering out low-level components in the preprocessing stage. (**B**) Spline interpolation of σ_ITD_, for the full curve (red) or by splitting it in two halves at 0 μs (green). (**C**) Forcing left/right symmetry between ITD statistics (example in the left refers to σ_ITD_). The panels depict results for the CIPIC HRTF database.

**Figure 4 supplement:**
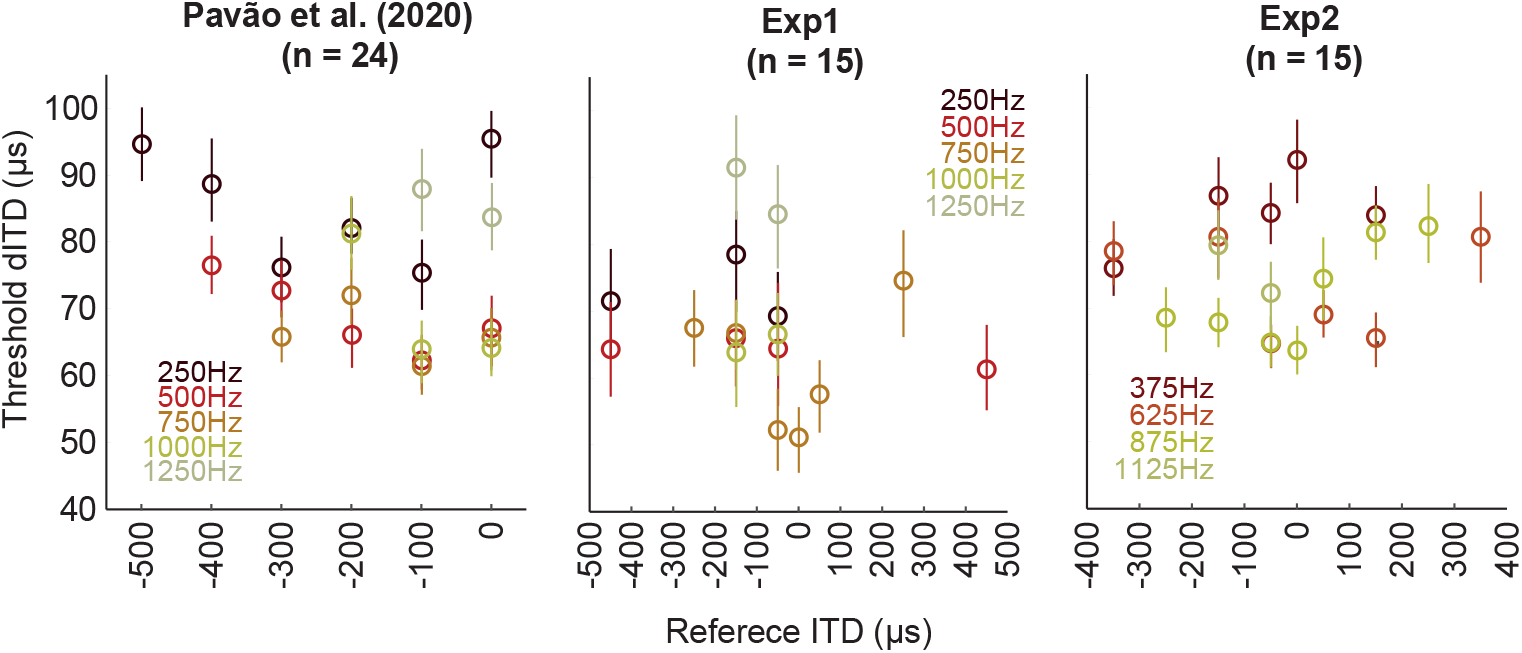
Average thresholds obtained for the ITD discrimination task. Mean dITD thresholds (CI = 50%) across conditions for each pool of subjects. Each experiment assesses different sets of reference ITD and frequencies. Notice that dITD thresholds are uncorrelated with either reference ITD or frequency.

### Prediction of dITD thresholds by √FI_ITD_ estimated from different methods

We used the linear mixed-effect models method (Magezi 2015) to assess the prediction of dITD threshold by √FI_ITD_. Each distribution of √FI_ITD_ constitutes a different model for predicting dITD thresholds. Each model relies on √FI_ITD_ estimates from a combination of the multiple methods, namely HRTF database (CIPIC or LISTEN), fraction of low amplitude components filtered out (from none to up to 50% percentile), ITD delay compensation (none or compensated), σ_ITD_ interpolation procedure (single interpolation across all azimuth range or two independent interpolations, across left azimuths and across right azimuths), and forced symmetry procedure (none or perfectly symmetric).

We computed the Akaike Information Criterion (AIC) for each model, and compared them using the simplified metric dAIC which indicates the distance to the best prediction model (**Figure 5A** and **Figure 5 supplement**). The best model was the CIPIC HRTF with forced symmetry (dAIC = 0); however there is no statistical evidence for selecting this best model against any other model for which dAIC < 2.

We found that 5 out of the 80 tested models were equivalently good for predicting dITD thresholds: all of them were based on the CIPIC HRTF and considered all low-level signal components (all depicted in **Figure 3** and **Figure 3 supplement**, with is based on the same dataset of **Figure 5A**). Four of them did not include ITD delay compensation: (a) direct method, (b) forced symmetry, (c) bilateral interpolation of σ_ITD_, (d) forced symmetry together with bilateral σ_ITD_ interpolation. The model with delay compensation and forced symmetry was also a good predictor of dITD thresholds: this was the sole model with ITD compensation out of the five best, and the one that got the dITD closer to the statistical threshold. Therefore, the best tradeoff between accurate dITD prediction and raw data manipulation is achieved by the √FI_ITD_ estimates using the direct method on the CIPIC HRTF database. **Figure 5B** shows normalized dITD thresholds vs √FI_ITD_ estimates for this model: the dispersion follows a power law with smaller dITD thresholds when √FI_ITD_ is high.

**Figure 5 supplement:**
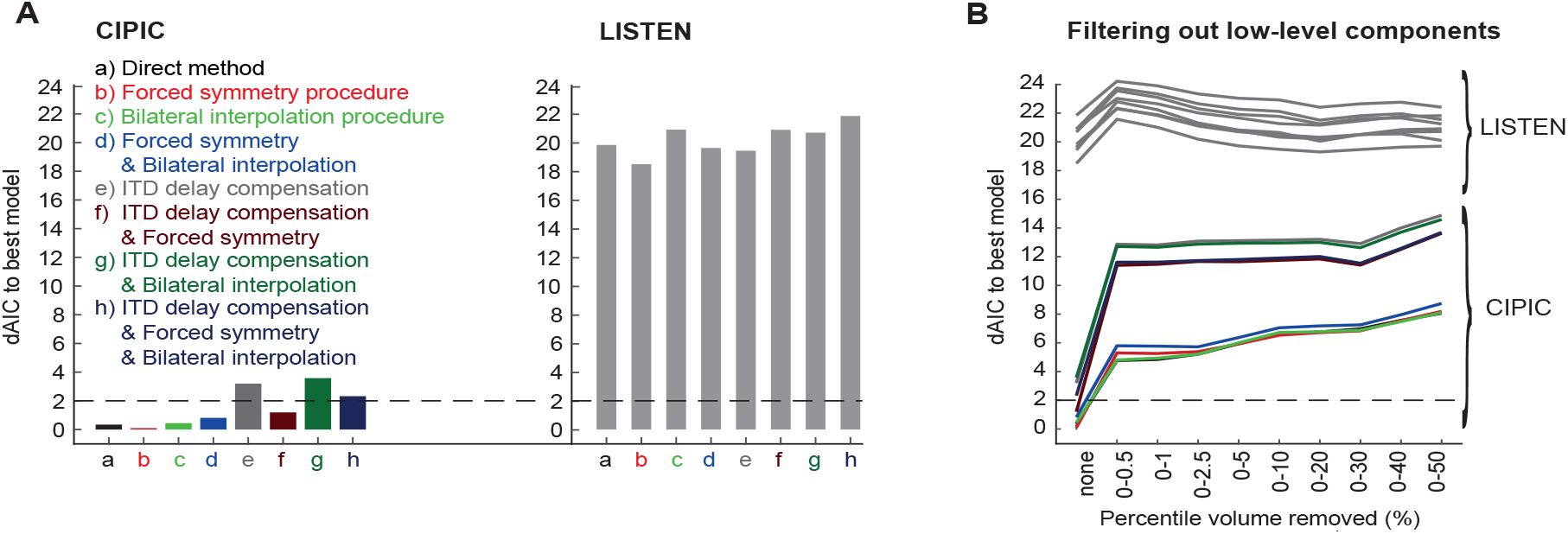
Prediction of dITD thresholds by the direct and additional methods for estimating √FI_ITD_. Models with dAIC values lower than 2 are considered to have the same predictive power. (**A**) Left panel shows models obtained using the CIPIC HRTF database; the right panel shows the ones obtained from the Listen HRTF database. (**B**) Low-level component removal in each of the methods shown in Figure 5A. Values of dAIC are lower when the percentage of low volume components removed is zero.

## DISCUSSION

The discrimination of sound positions in natural environments reflects the amount of azimuthal information carried by the stimulus, which can be measured from naturalistic sound statistics (Pavão et al. 2020). ITD Fisher Information √FI_ITD_ is an index that predicts ITD tuning in mammalian brainstem (Pecka et al. 2008; McAlpine, Jiang, and Palmer 2001), as well as human ITD discriminability for frequencies higher or equal to 500 Hz (Pavão et al. 2020). In other words, higher values of √FI_ITD_ related to more accurate measures of ITD discriminability, and vice-versa. This indicates that √FI_ITD_ could play a prominent role in human azimuth perception. However, √FI_ITD_ poorly predicted ITD thresholds at 250 Hz (Pavão et al. 2020). In the present study we considered different factors that could lead to this decreased performance, investigating methods that minimized noise on HRTF collection, mimicked auditory mechanisms, or reflected properties of natural environments, while trying to optimize the prediction of ITD thresholds from √FI_ITD_ at frequencies where this binaural cue is relevant. We found that direct √FI_ITD_ estimation from the CIPIC HRTF database is the best predictor of ITD discriminability, for reasons discussed in detail below.

### Prediction of dITD thresholds by √FI_ITD_ estimated using different methods

#### CIPIC vs LISTEN database

We found that estimating √FI_ITD_ from the CIPIC HRTF database resulted in an improvement of dITD threshold prediction when compared to estimates based on LISTEN HRTFs (**Figure 5A** and **Figure 5 supplement**). The LISTEN database provides HRIRs obtained in an anechoic chamber, differently from CIPIC. Moreover, CIPIC provides HRIRs with very short duration -- likely a strategy for avoiding echoes; however, the authors did not specify the features of the room in which those responses were obtained (Algazi et al. 2001).

We hypothesize that this outcome is related to the low signal-to-noise ratio in CIPIC HRTF signal collection, which better matches the power spectrum of natural sounds at lower frequencies (Młynarski and Jost 2014). In natural environments, reverberation is higher for low frequency sounds, increasing σ_ITD_ which, in turn, leads to reduced √FI_ITD_ (**Figure 2A**). This follows from out conjecture that ITD processing in the auditory system is driven by √FI_ITD_, thus requiring larger ITD differences for discriminating azimuths at lower frequencies. Therefore the CIPIC database, which has decreased √FI_ITD_ for the lowest frequencies, leads to better prediction of ITD discriminability.

#### Removal of low-level components

Only √FI_ITD_ estimates that considered the full range of low level audio components were good predictors of ITD discriminability (**Figure 5 supplement B**). In other words, models considering any form of filtering lead to higher dAIC, regardless of the HRTF database.

#### ITD delay compensation

One potentially important change for computing ITD statistics would be implementing internal delays in the √FI_ITD_ prediction model as suggested by Jeffress’ model (Joris and Yin 2007), a potential neural mechanism for ITD computation in brainstem. However, we found that delay compensation deteriorates prediction of ITD thresholds through all conditions (**Figure 5 supplement A**, left -- *a* vs *e, b* vs *f, c* vs *g* and *d* vs *h*). This supports alternative models for ITD coding, based on spike rate (Pecka et al. 2008; McAlpine, Jiang, and Palmer 2001) instead of labeled lines.

#### Interpolation on σ_ITD_

Although interpolating the FI curve through the full azimuth range generates an artificial smoothing for frontal azimuths, predictions employing this correction were slightly better when compared to those based on bilateral interpolation (i.e., on each side separately; **Figure 5 supplement A**, left -- contrast *b* with *d, e* with *g, f* with *h*). Thus, bilateral interpolation is likely to be more destructive than direct interpolation over the full range of azimuths, despite the artifacts introduced by the latter.

#### Forcing symmetry

Although human heads and ears exhibit bilateral symmetry, the estimated μ_ITD_ and σ_ITD_ were slightly asymmetric, suggesting minor systematic measurement errors. Enforcing these statistics to be symmetric by construction leads to a symmetric √FI_ITD_, which better predicts ITD discriminability in most conditions (**Figure 5 supplement A**, left -- compare *a* vs *b, e* vs *f, g* vs *h*; although *c* vs *d* shows the opposite result). Given that not all conditions including this modification lead to improved prediction, and that improvements due to forced symmetry compared to the direct method on CIPIC are not statistically distinguishable, Occam’s razor dictates that we should select the simpler method.

Our results further support that natural ITD statistics predict human ITD discriminability, even for sound frequencies below 250 Hz. Differently from previous studies, our results indicate that ITD statistics estimated from HRTFs with noisy low-frequency components better predict ITD discriminability thresholds than those computed from noiseless HRTF, as the former more closely approximates the properties of sound in a natural environment. Moreover, our results support that compensating for ITD delays and filtering out low-level signal components are detrimental to effective prediction of human ITD discriminability.

## MATERIALS AND METHODS

### HRTFs measurement

HRTF used in this study were extracted from the publicly-available LISTEN and CIPIC databases (http://recherche.ircam.fr/equipes/salles/listen and http://sofacoustics.org/data/database/cipic). The LISTEN HRTF database consists of HRIRs recorded at the Institute for Research and Coordination in Acoustics/Music (IRCAM). The measurement was performed in an anechoic room that absorbed sound waves above 75 Hz. We selected the HRIRs from frontal azimuths in elevation zero, resulting in 13 azimuths from −90º to +90º, with steps of 15º, from all the 51 subjects available.

The CIPIC HRTF database was made available by the U. C. Davis CIPIC Interface Laboratory (Algazi et al. 2001). We selected the HRIRs from frontal azimuths (−80º, −65º, −55º, from −45º to 45º in steps of 5º, at 55º, 65º and 80º) at zero elevation, from all the 45 subjects available.

### Estimation of ITD Fisher information

#### Direct method

The √FI_ITD_ estimation procedure described in (Pavão et al. 2020) is referred to as the “direct method”. We applied this method to both the LISTEN, as in (Pavão et al. 2020), and the CIPIC HRTF database. In **Figure 2**, we describe how FI_ITD_ can be estimated using the CIPIC database. We examined the full range of frequencies for which ITD is relevant for sound localization: 250 Hz to 1250 Hz, sampled with steps of 125 Hz. We analyzed the effect of a number of modifications of the direct method, both isolated and in combination. Each of these modifications is described in detail below, as well as outlined in **Figure 3** and **Figure 3 supplement**.

#### Removal of low-level components

The cochlea only starts to vibrate, and therefore excite hair cells, given a minimal sound pressure level which leads to intensity thresholds (Pickles 2008; Gelfand 2017). We computed √FI_ITD_ while removing waveform segments whose maximum amplitude was lower than a range of cutoffs (quantiles 0, 0.005, 0.01, 0.025, 0.05, 0.1, 0.2, 0.3, 0.4, and 0.5). A sample √FI_ITD_ obtained for a given threshold is shown in **Figure 3 supplement**.

#### ITD compensation

In classical models for binaural hearing, it is proposed that different ITDs excite distinct coincidence detectors after adequate compensation by internal axonal delay differences (Joris and Yin 2007; Stern and Colburn 1978). We expressed this concept in an alternative √FI_ITD_ estimation procedure, by subtracting the mean ITD for each azimuth, leading to distinct values of standard deviation (σ_ITD_). The resulting √FI_ITD_ is shown in **Figure 3B**.

#### Interpolation procedure on σ_ITD_

HRIR recordings consider a limited number of positions along the horizontal plane. The interpolation of σ_ITD_ across the complete range of azimuths, as applied in the direct estimation method, generates an artificial smoothing at frontal azimuths. Here, we interpolated σ_ITD_ across left azimuths independently of the right azimuths, thus obtaining a √FI_ITD_ curve with a sharp peak around zero (**Figure 3 supplement B**).

#### Forced symmetry procedure

The subject-averaged μ_ITD_ and σ_ITD_ estimated from HRIRs are slightly asymmetric, which suggests the existence of minor (though systematic) measurement errors. To mitigate this, we forced the symmetry of μ_ITD_ and σ_ITD_ curves, by averaging each statistic across actual and mirrored azimuths, leading to symmetric √FI_ITD_ (**Figure 3 supplement C**).

#### Estimation of dITD thresholds

In order to select the method for computing √FI_ITD_ that best predicts human ITD thresholds, we pooled the previously collected data for 24 subjects (12 females and 12 males) by (Pavão et al. 2020) with data for 30 subjects (9 females and 21 males) collected specifically for our study in two experimental phases. The protocol was approved by the Ethics Committee of Universidade Federal do ABC. There were no distinctions between groups used in both studies. The subjects were tested in different sets of combinations of frequency and reference ITD, as shown in **Table 1**.

**Table 1:**
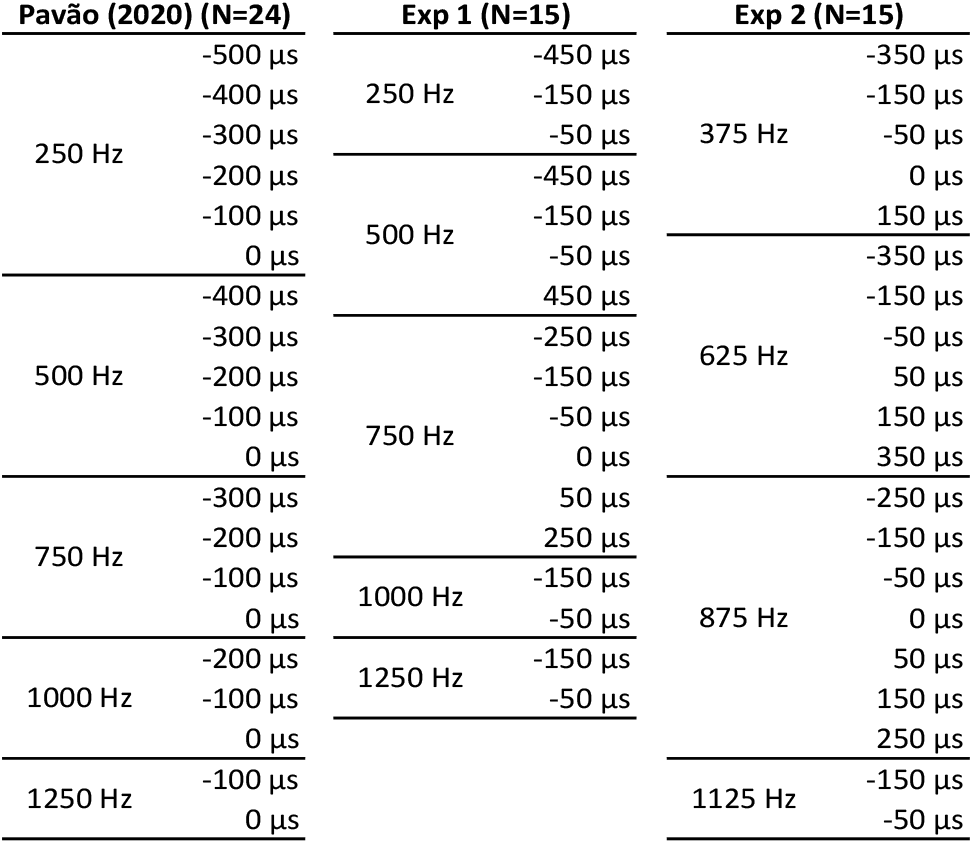
Subjects and experimental conditions in the ITD threshold protocol (FI_ITD_).

Each of these conditions consisted of 25 trials. In each trial, two sounds (10-element bursts) were presented in sequence. The first half of each sound defines the *reference* ITD, while the second half either repeats this reference or introduces a different ITD; the changes were always towards the periphery, in varying amounts. The subjects had to respond whether or not (or unsure) they perceived a change in the location of the second sound with respect to the first. A schematic trial describing the protocol is presented in **Figure 4**. We assigned the value 0 for incorrect and unsure responses, and 1 for correct responses; and estimated the ITD discriminability threshold (dITD) that led to maximum hits and minimum false positives, using a ROC classifier (Pavão et al. 2020). **Figure 4 supplement** shows dITD thresholds across conditions for each subject pool (mean and 50% confidence interval, computed using bootstrap as in Pavão et al. 2020).

#### Prediction of dITD thresholds by ITD Fisher information

We assessed the prediction of dITD threshold by each of the aforementioned methods for computing √FI_ITD_ using linear mixed-effect models (Magezi 2015), considering √FI_ITD_ of the reference ITD as a fixed factor and subject id as random factor. Since dITD threshold and √FI_ITD_ are both well described by power laws, we linearized them using a logarithmic transformation in both variables. We calculated the Akaike Information Criterion (AIC) for comparing each method (Pavão et al. 2020). The arbitrary scale of AIC was subtracted from the AIC of the best predictor, such that the best predictor corresponded to zero dAIC. Predictors with dAIC between 0 and 2 were considered equally good in power. This analysis is shown in **Figure 5** and **Figure 5 supplement**.

